# Complex interspecific interactions influence the interactions between pest control and pollination in coffee agroecosystems

**DOI:** 10.1101/2024.01.12.575383

**Authors:** Chatura Vaidya, Gabriel Dominguez Martine, John Vandermeer

## Abstract

Ecosystem services mediated by biodiversity are essential for the well-being of human beings. While there is ample research on individual ecosystem services (such as pollination, nutrient cycling), there is now growing recognition to examine the interactions between multiple ecosystem services and their contribution to productivity in order to manage agroecosystems sustainably. In this study, we examined the interactions between pollination and pest control in coffee agroecosystems in Chiapas, Mexico. We tested how management of shade trees, particularly of nitrogen-fixing shade trees, at the farm scale mediated the outcome of the interactions between two ES. We found that there was no trade-off between pest control and pollination services despite the deterrence of pollinators by the dominant and aggressive ant species, *Azteca sericeasur*, which also controls the coffee berry borer, a major pest of *Coffea arabica*. We found additive effects of pest-control and pollination on early fruit set and fruit weight of coffee plants. Proximity to nitrogen-fixing shade trees had indirect effects on pest-control via the reduction of *Azteca sericeasur* activity on the coffee bushes. These findings highlight that ecosystem services are a result of complex interspecific interactions and that biodiversity-friendly management practices can promote favorable outcomes of these interactions on coffee yield.

## Introduction

Provisioning services such as food, timber, and fiber are essential to human well-being, and much research has focused on maximizing their availability by improving the management of ecosystems. Management practices can also influence the ecosystem services (hereafter, ES) that support these provisioning services, including pest control, pollination, regulation of soil fertility and nutrient cycling. Not surprisingly, a majority of the research has considered each of these ES in isolation, perhaps necessary so as to navigate around the complexity that arises, focusing on a single ES such as pest control or pollination. There is now a growing recognition that multiple ES interact with one other and that such complexity must be addressed in search of sustainable ecosystem management and the promotion of biodiversity(Chain-guadarrama et al., 2019).

Animals, including many insect species, are important pollinators for about 88% of angiosperm species, providing vital ecosystem services for wild plants and food crops worldwide (Ollerton et al., 2011). Similarly, pest control, another animal-mediated ES increases crop productivity by reducing pest damage to crops (Gutierrez-Arellano and Mulligan, 2018). Both these services have significant economic importance and therefore have received considerable attention. In the past decade, a literature has evolved focusing on the interactions between pollination and pest-control in different agroecosystems. Most of these studies have found varying effects of the interactions between these two ES on crop yields (Garibaldi et al., 2018).

While some studies have found synergistic effects (the gain in crop yields when pest control and pollination interact is higher than the sum of the gain when these two ES act separately) for example in red clover and oilseed rape (Albrecht and Sutter, 2016; Lundin et al., 2012), others have found that they only act independently (no interaction) for example in cucumber crops (Barber et al., 2012), one study has found a weak negative interaction on oil seed rape yields (Bartomeus et al., 2015). Thus, even though the interactions between these two ES have variable outcomes, there is evidence that these services complement or augment one another.

Farmers worldwide, augment soil fertility by adding fertilizers – organic or synthetic, or by using cover crops or nitrogen fixing shade trees with the direct goal of supplying limiting nutrients to the plants. Nutrient availability not only affects plant productivity but can also have bottom-up effects and potentially influence a variety of interactions between trophic levels. Nitrogen (N) can influence plant-pollinator interactions through altered floral phenology (Cleland et al., 2006; Hoover et al., 2012), changes in floral production (Burkle and Irwin, 2010) and floral rewards such as nectar volume, nectar composition and pollen composition (Ceulemans et al., 2017; Gardener and Gillman, 2001). Similarly, N may also limit herbivorous insects and N availability can have direct significant effects on herbivore development, growth and reproduction and potentially increase mutualistic interactions between honeydew-producing insects and ants (Gonthier et al., 2013; Vaidya and Vandermeer, 2021). One way that soil fertility is enhanced in tropical agroecosystems is by incorporating N-fixing trees. Several studies have reported an increase in N in crops when associated with N-fixing leguminous trees, either through direct N transfer via mycelial networks or tree root exudate absorption by crops or through prunings via leaf litter mulching (reviewed in Nygren et al., 2012).

Coffee agroecosystems are excellent model systems for the study of interactions between different ES, since coffee is grown on 11million hectares of land in the tropics by millions of farmers worldwide (FAO, 2015), its importance both environmentally and economically is large. It is currently grown under a variety of management conditions from “rustic coffee” to “sun coffee” (Moguel and Toledo, 1999), and farms that grow coffee under the canopy of shade trees have been shown to support higher associated biodiversity and the provisioning of multiple ES such as pollination, pest control, carbon sequestration, and enhanced nutrient cycling (Greenberg et al., 1997a, 1997b; Jha et al., 2014; Perfecto et al., 1996; Perfecto and Vandermeer, 2008; Philpott et al., 2006; Tscharntke et al., 2011), thus making coffee agroecosystems ideal for studying the interactions among ES. Interactions between ES can either be direct – via the interactions of the species that provide the different ES, or indirect – via changes in a shared driver (Bennett et al., 2009). In the coffee plantations of southern Mexico, farmers regularly grow coffee plants under the canopy of nitrogen-fixing shade trees from the genus *Inga* (Perfecto and Vandermeer, 2002) and there is some evidence of N transfer to coffee plants from the *Inga* trees (Grossman et al., 2006; Roskoski, 1982). Thus, inquiring if this shared driver, nutrient availability, modifies the interaction outcome between pest control and pollination via indirect effects is an important piece of the puzzle.

Most research on the interactions between pest-control and pollination in coffee agroecosystems has focused on pest-control by birds and has found that these interactions have variable outcomes, from synergistic (Martínez-Salinas et al., 2022) to complementary(Classen et al., 2014). But there are several other kinds of organisms that provide pest control and among them are ants (Anjos et al., 2022). Ants provide pest-control in coffee agroecosystems (Jha and Vandermeer, 2010) and the mechanisms through which they control pests are many. For example, ant-hemipteran mutualistic interactions are abundant in nature (Delabie, 2001) in which ants protect their mutualistic partners from their predators and in doing so, also protect plants from other potentially more damaging pests (Floate and Whitham, 1994; Perfecto and Vandermeer, 2006).

However, another consequence of the ant-hemipteran mutualism, especially in aggressive ants, is that ants also seem to “defend” plants from all plant visitors, including pollinators. Ants can chase away pollinators (Philpott et al., 2006; Vannette et al., 2017) and indirectly deter pollinator visitation by stealing nectar from the flowers (Ghazoul, 2001; Levan and Holway, 2015) and leaving pheromones on flowers that pollinators then tend to avoid (Lach, 2007; LeVan et al., 2014; Sidhu and Rankin, 2016). Thus, ant-hemipteran mutualisms may benefit the plant from defense against other herbivores yet may simultaneously reduce the effectiveness of the plant-pollinator mutualism thus, resulting in an apparent tradeoff (Fig 1).

**Figure 1.**
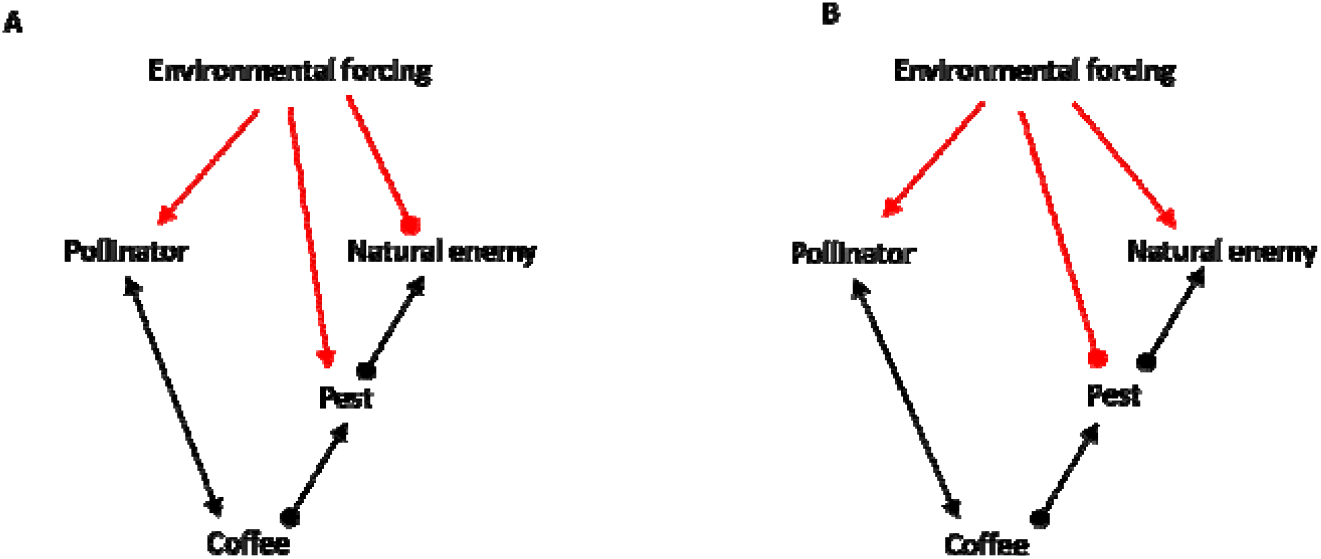
Conceptual framework showing two scenarios. Arrows show a positive effect and filled circles show a negative effect. A) depicts a trade-off between pollination and pest control when environmental conditions have a positive effect on pollinators and pests but have a negative effect on the natural enemies of the pest. B) depicts a potential positive or no interaction between pollination and pest control when environmental conditions have a negative effect on the pest and a positive effect on pollinators and the natural enemies of the pest. There can be several such permutations that may give rise to trade-offs or synergistic interactions between pest control and pollination services.

In this study, we tested the interactions between two key ES -pest control and pollination via the interactions between ants and pollinators of *C. arabica*. Specifically, we investigated 1) whether the interactions between pest control and pollination are synergistic or concessionary (i.e., represent a trade-off) and, 2) whether management factors, particularly the association with nitrogen-fixing shade trees mediates the outcome of these interactions on coffee yield (Fig 1)

## Methods

### Study System

*Coffea arabica* is self-compatible but can benefit from animal pollination via increased fruit quantity and quality (Klein et al., 2003a; Philpott et al., 2006; Ricketts et al., 2004). Coffee (*Coffea arabica*) flowers in the dry season between February-May, the flowers remaining open for ∼2 days or, if unpollinated, for as much as 5 days. Its berries are harvested typically between September-December. if unpollinated, can remain open for ∼5 days. Coffee is frequently planted under the canopy of shade tree species. On Finca Irlanda, the site of our study, shade trees of approximately 90 different species are planted, with the most common ones fixing nitrogen and belonging to the genus *Inga* (Schmitt and Perfecto, 2021). The second most common shade tree is *Alchornea latifolia*. Since the farm is an organic plantation, no pesticides or herbicides are used, and manual weeding is carried out using machetes. *Coffea arabica* has numerous pests, but the coffee berry borer (CBB, *Hypothenemus hampei*) and the green coffee scale (GCS, *Coccus viridis*) are considered to be the most prominent pest species. The CBB female bores and lays eggs inside the coffee berries and the larvae develop by eating the pulp of the berry, causing extensive damage to it. The GCS is a phloem feeder and forms mutualistic interactions with several ant species. In our study site, their most prominent interactions are with the dominant ant species, *Azteca sericeasur* (Perfecto and Vandermeer, 2006). These ants defend the GCS aggressively from its predators and other competing herbivores thus indirectly controlling the coffee plants from CBB infestation (Morris et al., 2015; Vandermeer et al., 2010). *Azteca sericeasur* does not discriminate between the natural enemies of GCS and other organisms and is known to be aggressive towards any organism that visits the coffee plants, including pollinators and other ant species (Philpott et al., 2006; Vandermeer et al., 2010; Vannette et al., 2017). Additionally, *A. sericeasur* nest arboreally nesting inside the tree trunks and occasionally making large carton nests on the trunk surface. They forage extensively on neighboring coffee bushes, mostly via tending the GCS on these bushes. The GCS reach their highest densities on coffee bushes closest (within 5 m) to those shade trees that house nests of *A. sericeasur* ants (Perfecto and Vandermeer, 2006; 2019).

### Study Design

This study was conducted in Finca Irlanda from February 2018 to December 2019. Finca Irlanda is a 300 ha organic shaded coffee plantation in Chiapas, Mexico (92°20′29′′□W and 15°10′6′′□JN) at an elevation range between 900 and 1200 m a.s.l, with mean annual rainfall of approximately 4500 mm (Schmitt and Perfecto, 2021).

We selected coffee bushes within 2m of shade trees; shade trees were either nitrogen-fixing shade trees from the genus *Inga* (n=33 in 2018 & n=23 in 2019) or non-nitrogen fixing shade trees (non-*Inga spp*., multiple species-most commonly *Alchornea latifolia*, n=34 in 2018 & n=28 in 2019). Roughly half of the shade trees had *A. sericeasur* (hereafter, Azteca ants) nests on them (Fig 2) and foraging on coffee bushes near the shade trees, while the other half did not. Thus we had a paired design – coffee bushes paired with *Inga* spp. with Azteca ants and without ants, coffee bushes paired with non-*Inga* spp. with Azteca ants and without ants.

**Figure 2.**
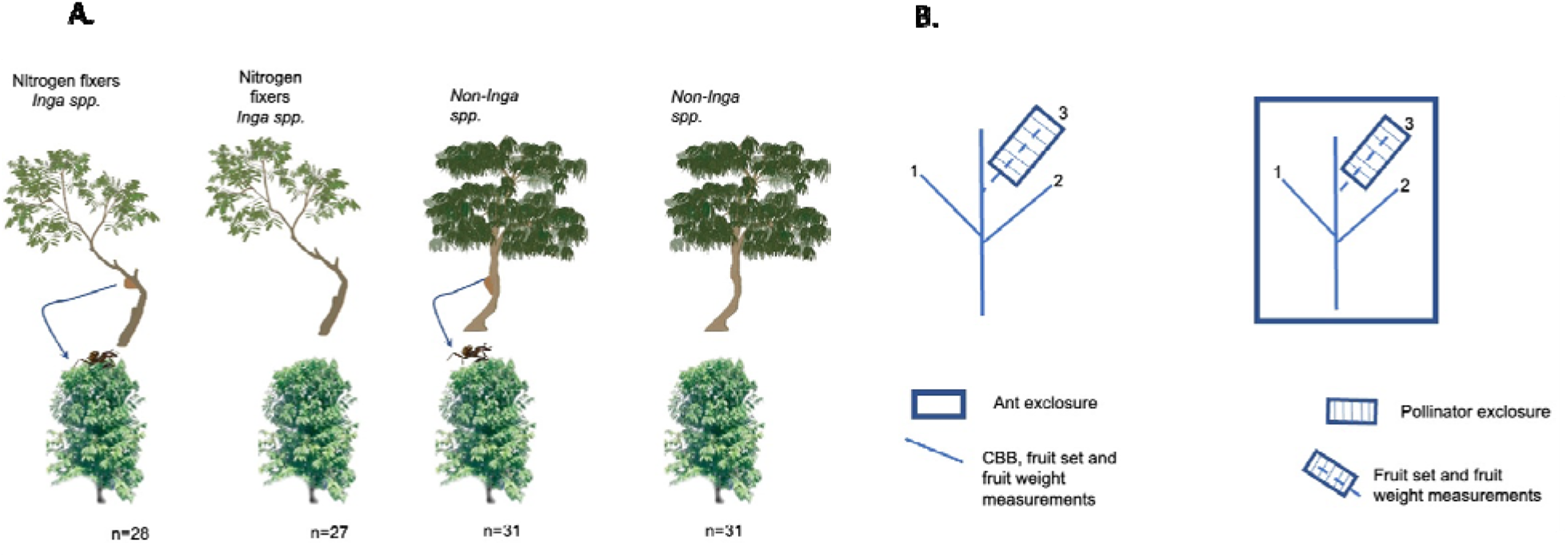
Experimental design. A) Full-factorial design of shade tree (N fixation), ant exclosure and bee exclosure treatments. We selected coffee bushes paired with *Inga spp*. with *Azteca seriseasur* and without any ants and those paired with non-*Inga spp*.(non-N fixers) with *Azteca seriseasur* and without any ants. B) On the plant level, two branches were kept open to pollination and one branch was excluded from ants and both pollinators (bagged). We assesed fruit set and fruit weight on all three branches, while CBB presence/absence in fruits was only assesed on branches that were open to pollinators (1,2).

On each coffee bush, we selected 3 branches, two of them were kept open to animal pollinators (open pollination) and one branch was bagged (0.8-1mm mesh size) to exclude animal pollinators (bagged pollination). On each coffee bush, before coffee started flowering, we counted the total number of buds on all three branches in February 2018 and 2019. On coffee bushes that did not have Azteca ants, we applied tanglefoot at the base of the branch to exclude other ant species from all three branches (ant exclosure), while on bushes that had Azteca ants, tanglefoot was applied only on the bagged branch. Bags were removed at the end of the flowering period (end of April). Thus, we had three factors in total – pollination (open and bagged), ants (with Azteca ants and without ants) and Shade treatment (*Inga spp*. and non-*Inga spp*.) We measured Azteca ant activity on the coffee bushes by selecting a random point on the plant and counting the total number of ants passing the point for a period of one minute. Azteca ant activity ranged from 5 – 20 ants per minute.

#### Pollinator visitation

When coffee plants started flowering, we counted the number of open flowers on the open pollination branches on each bush and during 10 min timed observations recorded the species of the visiting pollinators, the total number of flowers visited by the pollinators and their visit duration. We pooled total number of flowers and visits from the two open branches for each coffee bush and calculated visitation rates by dividing the number of visits by the total number of flowers.

#### Fruit quality and quantity

We counted the total number of fruits at two time points – once in May-June, accounting for initial pollination, and during harvest, accounting for late pollination. We counted initial and late fruit set by dividing the total number of flower buds by the total number of berries during each count respectively for all branches (open and bagged). For branches that were open to pollination, we averaged fruit set across the two branches. We harvested all available berries on the three branches of each bush and measured the weight and counted the number of beans in each harvested berry. Since the presence of one bean in a berry or fruit (peaberry) is a sign of unsuccessful pollination, this measure was useful to determine pollination success rates among the different control and treatment groups. Fruit set on bagged branches was only counted in 2019, but fruit weight and the number of beans in each fruit was measured in both years.

#### Coffee berry borer control

We recorded the presence/absence of the CBB in each harvested berry from all the branches that were open to pollination. Since the bagged branches on coffee bushes with Azteca ants had tanglefoot applied on them, we only compared branches open to pollination for the control of CBB.

#### Effect of shade trees

Along with total number of flowers on the three branches, we also measured the corolla diameter of five flowers on one branch of all coffee bushes to understand the effect of N availability on floral traits. We sampled five leaves each from the shade trees (8 *Inga spp*. and 8 non-*Inga spp*.) and their paired coffee bushes (nine of each). We ground dried samples using a coffee grinder and used a representative sub-sample to analyze % C and % N using a LECO Trumac CN combustion analyzer (LECO Corporation, St. Joseph, MI). We used the total C and total N data to calculate the carbon to nitrogen ratio (C:N).

### Statistical Analysis

To understand if pollinator visitation differed between the treatments, we constructed a generalized linear mixed-effects model with a zero-inflated negative binomial distribution using the package “glmmTMB”, with number of visits by pollinators as the response variable and log(number of flowers) included as an offset. Predictor variables were the ants and shade tree treatments as well as year to account for interannual variation in pollinator visits as fixed effects and tree and site as random effects. We set the zero-inflation formula equal to 1, to denote equal probability of producing zeros over all observations. To determine whether the presence of the green coffee scale insects influenced ant visitation to coffee flowers, we performed a Fisher’s exact test.

We analyzed the effect of the three factors on both the initial fruitset and fruitset at the time of harvest by constructing linear mixed effects models using the ‘lmer()’ function from ‘lmerTest’. We constructed models with all three treatments-pollination, Azteca ants and shade trees-including two-way and three-way interactions and year as fixed effects. In both fruitset (initial and at the time of harvest) models, plant and site were added as random effects. Models with interactions were compared with those without interactions using likelihood ratio tests and AIC criteria using the “anova” function and the model with the lowest AIC score was chosen. To meet conditions of normality, fruitset at the time of harvest was arcsine square root transformed. We checked for model compliance with assumptions using the ‘DHARMa’ package.

To determine the effect of all treatments and their interactions on the weight of the coffee fruits we constructed linear mixed effects models; year was also added as a fixed effect. Branch, plant and site were added as random effects. Models with interactions were compared with those without interactions using likelihood ratio tests and AIC criteria using the “anova” function and the model with the lowest AIC score was chosen. To evaluate the incidence of peaberry formation, we constructed a generalized linear mixed effects model with a binomial error distribution with all three treatments and their interactions and year as fixed effects and plant as random effect. We constructed models with and without interactions and checked goodness of fit using AIC information.

We observed a decrease in ant activity in 2019 from the previous year, therefore we tested if this difference was statistically significant by fitting a generalized linear mixed effects model with poisson log link function. We added shade trees, year and their interactions as fixed effects and plant and site as random effects for this model. To account for the inter-annual variation in ant activity, we tested the effect of treatments on CBB status within the coffee berries (presence, absence) for the two years separately. We constructed generalized linear mixed effects models with a binomial error distribution and modeled the log odds ratio of CBB presence in the berry, CBB absence in the berry as the response variable and shade tree treatment, ants treatment and their interaction as fixed effects; plant and site were added as random effects. Models with and without interactions were tested for goodness of fit using AIC information.

For all the mixed effect models described above, we performed Type II Wald chisquare tests on the selected model to determine which effects were significant. We then used post-hoc tests to estimate marginal means and contrasts to make pairwise comparisons between the treatment levels if they were significant using the “emmeans function” in the “emmeans” package.

To test the effect of shade trees on the number of flowers, we constructed a generalized linear mixed-effects model with poisson log-link function. Shade trees and year were added as fixed effects and plant and site as random effects. To test the effect of shade trees on the size of the flowers, we constructed a linear regression model with shade trees and year as the fixed effects and checked if the model satisfied conditions of normality by performing a Shapiro-Wilk test on the residuals.

To test whether there were any differences in %N and C:N ratios between *Inga spp*. and non-*Inga spp* shade trees and between the paired coffee bushes, we conducted a Wilcoxon test. All data was analyzed using R version 4.1.2.

## Results

### Pollinator visitation

There were a total of 1149 visits to coffee flowers in both years by pollinators; 87% of the visits were made by the introduced *Apis mellifera*, 12.7% of visits were made by native stingless bee species *Scaptotrigona mexicana, Trigona fuliventris* and *Trigona nigerrima* and the remaining visits were by the bee species *Agapostemon sp*.

Bee visits to coffee flowers on bushes with Azteca ants were significantly lower (3.8 ± 8.1; mean ± std. dev.) than on those with ant exclosures (15.6 ± 15.4, χ2 = 21.5, d.f. = 1, p < 0.001, Fig 1A) and were higher in 2019 (χ2 = 12.6, d.f. = 1, p < 0.001). We did not find significant effects of shade treatment on bee visitation. Additionally, bee visit duration was strongly reduced on coffee bushes with Azteca ants than without ants (Fig 1B).

Additionally, we found that Azteca ants tended to forage in the coffee flowers significantly more in the absence of the GCS (p=0.02).

### Fruit set

Early fruit set significantly increased from 0.45 (bagged pollination) to 0.65 (open pollination) with pollinator activity (χ2 = 27, d.f. = 1, p < 0.001 Fig 4). Additionally, early fruit set also increased significantly from 0.5 to 0.6 when Azteca ants were present on coffee bushes (χ2 = 9.9, d.f. = 1, p= 0.002). The selected model with lowest AIC score did not have three-way or two-way interactions as fixed effects and shade tree treatment was not significant. Fruit set was also significantly higher in 2019 than in 2018 (χ2 = 9.6, d.f. = 1, p =0.002).

**Figure 4.**
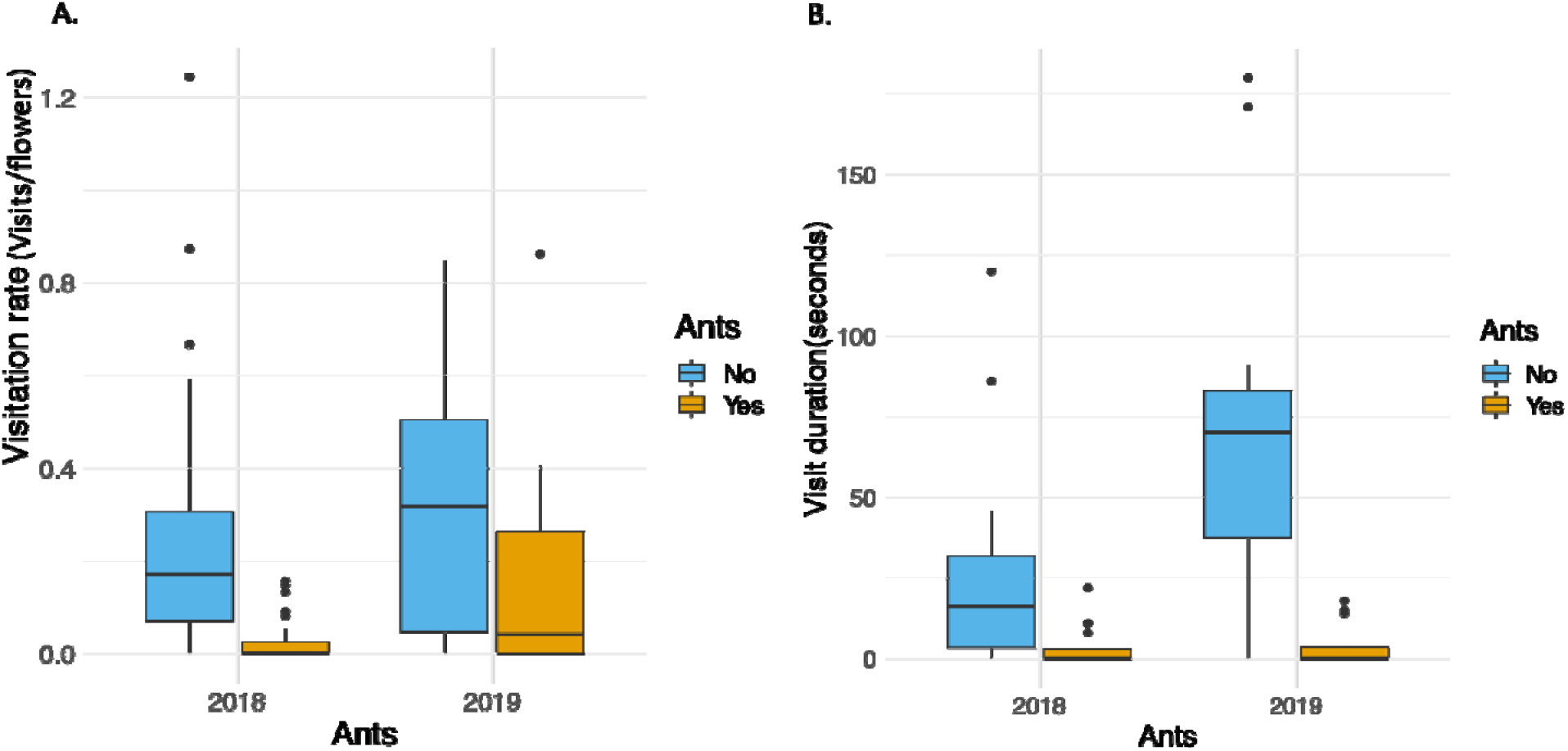
Visitation rate (A) and visit duration (B) of pollinators was significantly lower in the presence of Azteca ants. Both visitation rate and visit duration was higher in 2019.

**Figure 4.**
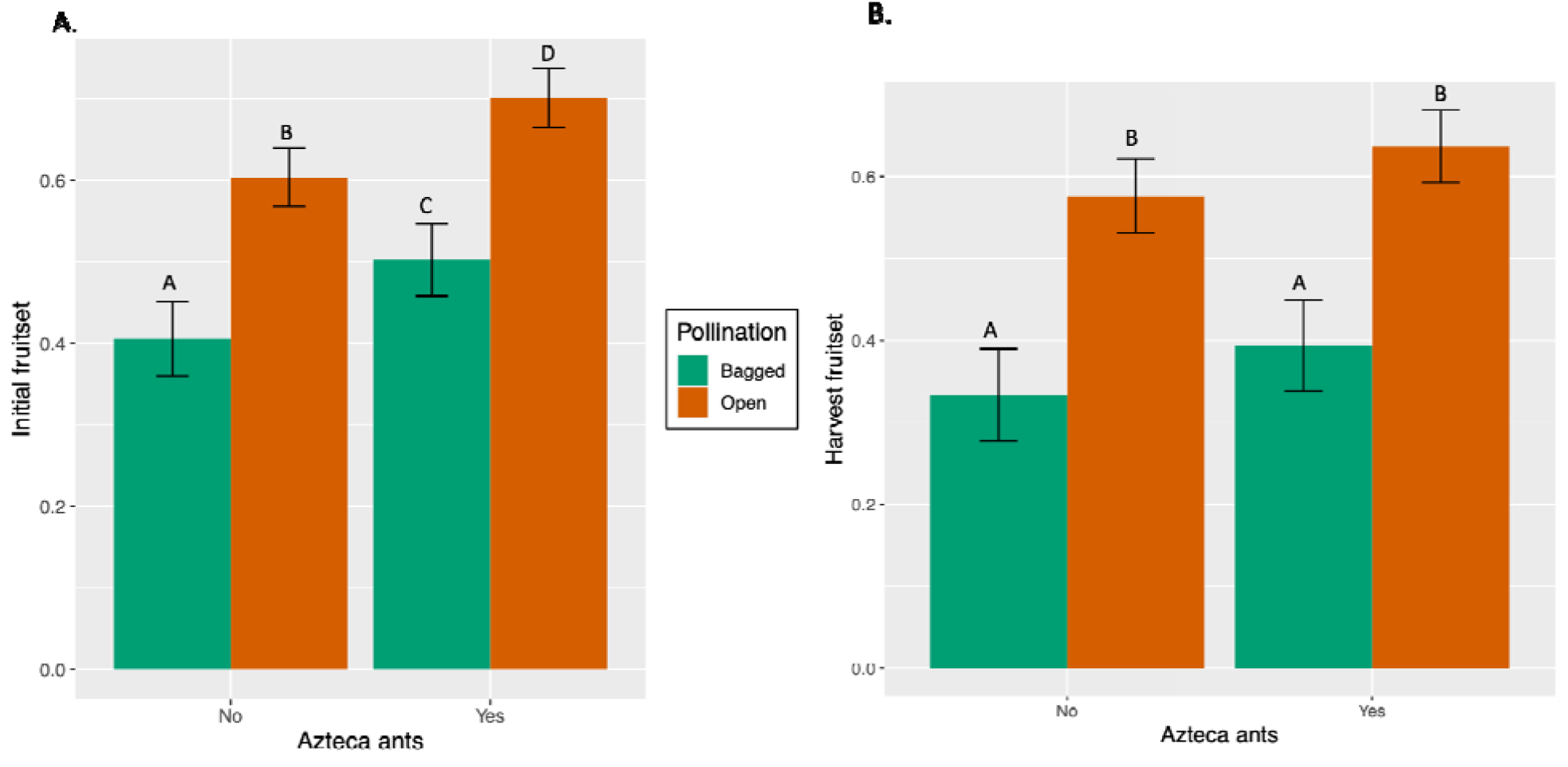
Effects of pollination and ant treatments on the intiial fruit set and fruitset at the time of harvest. Estimated marginal means with SE of the A) initial fruitset and B) fruitset at the time of harvest. Pollination treatments were significant for both initial and harvest fruit set (denoted by different colored bars in A and B). Presence of Azteca ants significantly increased fruitset in both bagged and open pollination treatments for the initial fruit set (A) but this effect was no longer significant for the fruit set at the time of harvest. Different letters denote statistically significant differences.

For fruit set at the time of harvest, the model with only pollination and year was the best fit model (ΔAIC = -5.1 for model omitting two-way and three-way interactions and all other fixed effects). Fruit set at the time of harvest significantly increased from 0.37±0.05 to 0.61± 0.03 when pollinators were allowed access (χ2 = 27.6, d.f. = 1, p < 0.001). There was no significant difference between the two years.

### Fruit quality

The best model for fruit weight was the one without two-way and three-way interactions. Fruit weight increased significantly from 1.56 g ± 0.04 to 1.74 g ± 0.03 (estimated mean ± std error) with pollinator activity alone representing an 11.5% increase (χ^2^ = 29.07, d.f. = 1, p < 0.001, Fig 5) and by 0.07g and in the presence of Azteca ant activity (χ^2^ = 4.23, d.f. = 1, p =0.039) representing a 4.3% increase in fruit weight.

**Fig 5.**
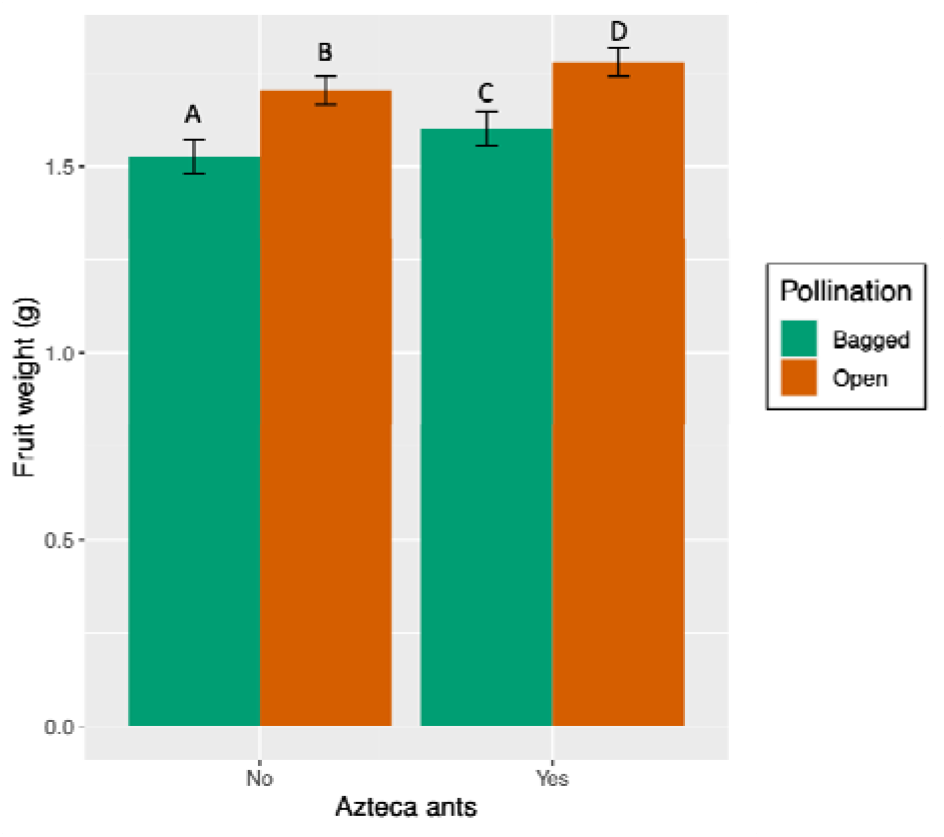
Effects of pollination and ant treatments on the weight of the coffee fruits. Estimated marginal means with SE were significantly different in the bagged and open pollination treatments, denoted by colored bars. Presence of Azteca ants significantly increased fruit weight in both bagged and open pollination treatments. Different letters denote statistically significant differences.

Probability of peaberry formation was significantly lower when pollinators were allowed access than on those with pollinator exclosures (Z = -2.199, p = 0.028). The model with only pollinator exclosure as the fixed effect was the best fit model (ΔAIC = -5.1 for model omitting two-way and three-way interactions and all other fixed effects).

### Pest control of CBB

We found that ant activity was significantly reduced in 2019 (χ^2^ = 58.2, d.f. = 1, p <0.001). On coffee bushes paired with non-*Inga* spp, Azteca ant activity was significantly higher in 2018 than 2019 (Tukey test, ß = 1.28±0.13, z = 9.53, p<0.0001). We therefore decided to look at CBB control for the two years separately.

For the year 2018, the interaction effect of the shade treatment and pest-control treatment was the only significant effect (χ^2^ = 4.2, d.f. = 1, p =0.04). Probability of the presence of CBB in coffee berries was significantly lower in bushes with Azteca ants paired with non-*Inga* spp. than without Azteca ants. There was no effect of Azteca ants on the probability of the presence of CBB in coffee berries on bushes paired with *Inga* spp (Fig 6A).

**Fig 6.**
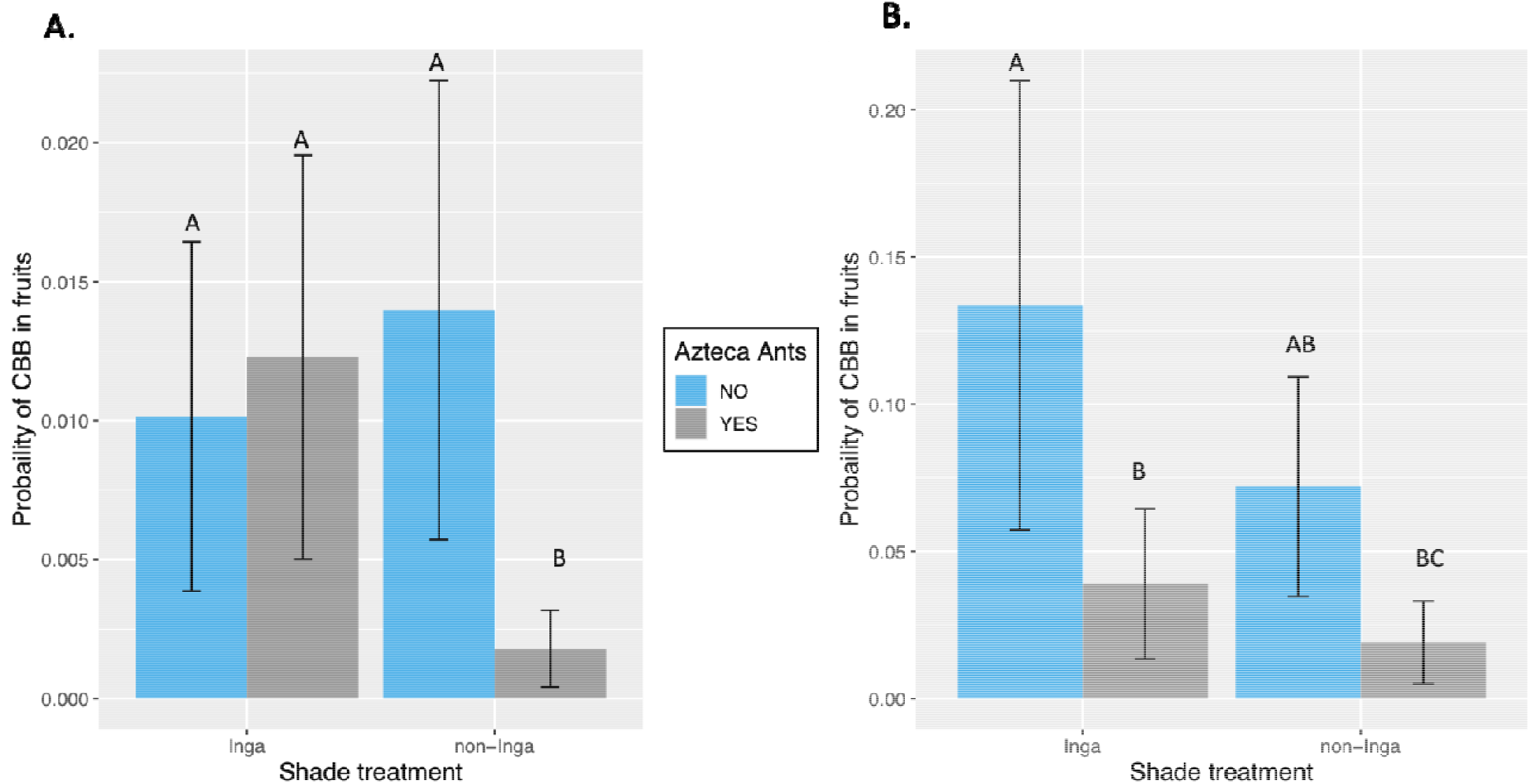
Estimated log odds ratio (probability) with SE of finding berries with CBB in them A) with and without Azteca ants on coffee bushes paired with *Inga spp*. and non-*Inga spp* in 2018, B) with and without Azteca ants on coffee bushes paired with *Inga spp*. and non-*Inga spp* in 2019. Different letters denote statistically significant differences.

For the year 2019, effect of Azteca ants alone was significant (χ^2^ = 5.1, d.f. = 1, p =0.02), with the probability of the finding CBB in coffee berries significantly lower on bushes with Azteca ants than on bushes with ant exclosures. Shade and ant treatments interaction did not have a significant effect on CBB control (Fig 6B).

## Discussion

Our findings show that there is no trade-off between pest control and pollination services despite the deterrence of pollinators by the dominant and aggressive ant species, *Azteca sericeasur*, that also controls the coffee berry borer, a major pest of *Coffea arabica*. Indeed, we found that there is an additive effect of pest-control and pollination on early fruit set and fruit weight. Proximity to nitrogen-fixing shade trees had indirect effects on pest-control via the reduction in Azteca ant activity on the coffee bushes, the reasons for which we discuss below.

Early and late fruitset both increased significantly when pollinators were allowed access to the plants (Fig4). This finding is in line with several studies that have found that although coffee is self-compatible, it benefits from animal pollination (Klein et al., 2003; Ricketts et al., 2004;

Martínez-Salinas et al., 2022). Azteca ants increased the benefit of animal pollination in early fruitset, which is surprising given that Azteca ants reduced both the number of visits and the visit duration of the pollinators visiting coffee flowers. Bees generally avoided visiting flowers on plants with Azteca ants to avoid aggressive interactions with the ants. Indeed, in some instances we noticed that Azteca ants would chase away bees attempting to visit coffee flowers. This phenomenon is common in interactions between aggressive ants and pollinators (LeVan et al., 2014; Sidhu and Rankin, 2016). While ants usually display this kind of aggressive behavior to protect their hemipteran partners, we found this behavior to be present even when the green coffee scale insects were absent on the coffee bushes. Bees likely also reduced their visits or visit duration due to resource competition with Azteca ants. We found that Azteca ants were foraging on coffee flowers, especially in the absence of the GCS, their hemipteran partners. This is contrary to past research that showed that ant-aphid mutualism increased ant floral visitation, reducing pollinator visitation and seed set(Levan and Holway, 2015). It is likely that they were foraging for nectar in coffee flowers to maintain their colonies in the absence of GCS, their major carbohydrate source. Thus, Azteca ants may have increased the initial fruit set of coffee via 2 non-exclusive mechanisms. First, since initial fruit set was significantly higher on both bagged branches and those open to animal pollinators, it is possible that Azteca ants were controlling floral and other herbivores which may have prevented resources being allocated from reproductive organs to vegetative tissues. Secondly, by foraging for nectar, ants may have increased self-fertilization in flowers in the open pollination treatments and may have also increased the pollination effectiveness of bees by reducing their visit duration to the flowers (Fig2B) and thereby facilitating outcrossing. The majority of the visits to coffee flowers were made by *Apis mellifera sculleta*, and previous studies have shown that interspecific competition for resources alters the behavior of honeybees and increases their pollination effectiveness by increasing the proportion of movement between trees (Brittain et al., 2013). Both these mechanisms seem likely and may have acted in concert to increase the initial fruit set. To disentangle these mechanisms, future research should compare the levels of floral and foliar herbivory on plants with and without Azteca ants and record fruit set. Observations of the movement of honeybees to plants with Azteca ants and the ones adjacent to them should either be made directly (Brittain et al., 2013; Greenleaf and Kremen, 2006) or by using fluorescent dye to detect pollen flow between them (Fitch and Vaidya, 2021). Additionally, flowers visited exclusively by Azteca ants should be marked to evaluate the overall effectiveness of ants as pollinators of coffee plants. Ants have been shown to be effective pollinators in certain plant species (Gómez, 2000); therefore their ability to pollinate coffee cannot be entirely ruled out. The effect of ants on the fruit set at the time of harvest was no longer significant (Fig 4B), and it is possible that plants aborted the fruits that were pollinated by Azteca ants (Rostás and Tautz, 2010). Nonetheless, fruit set was still higher on plants with Azteca ants suggesting that neither of these mechanisms can be discounted.

Fruit weight, like initial fruit set, also benefitted from pollinator and ant activity and increased by 15% in the presence of pollinators and Azteca ants. Other studies have found that pollinators, mainly bees, contribute between a 7-27% increase in fruit weights (Classen et al., 2014). Pollinators contribute to higher fruit weights by way of cross pollination which can decrease the likelihood of misshapen fruits, or in the case of coffee, peaberry (fruits with only one bean) formation (Boreux et al., 2013; Krishnan et al., 2012; Ricketts et al., 2004). In our study, access to pollinators reduced the probability of peaberries in coffee plants. Fruit weight increases with the number of beans by ∼0.6g (Supplementary info, Table 1), suggesting that peaberries reduce weights of the fruit by 0.6g, which is a substantial reduction in weight. Azteca ants likely contributed to the increase in fruit weight by 1) promoting movement of bees between plants and facilitating cross pollination as discussed above and 2) by controlling the CBB (Philpott et al., 2006)(Fig 6), which can significantly reduce fruit weights by 7.5% (Supplementary info, Table 1).

Ants controlled the CBB differentially in the two years. In 2018, we found that the interaction between Azteca ants and shade treatment alone was significant; no other effects were significant. The probability of finding CBB in coffee fruits was significantly lower only on plants paired with non-*Inga* spp. compared to the plants with ant exclosures (Fig 6A). In 2018, mean ant activity on plants paired with non-*Inga spp*. was higher than those paired with *Inga* spp. (mean ± std.dev, Non-*Inga spp*.17.1± 4.7; *Inga spp*.13.5± 5.8). *Inga* spp. trees usually have their own scale insects (*Octolecanium sp*.) that Azteca ants prefer to tend over the GCS. It is hypothesized that owing to the nitrogen fixation of *Inga* spp., honeydew quality or scale insect density is higher on Inga than the ones on coffee plants (Livingston et al., 2008; Vaidya and Vandermeer, 2021). Therefore, ant activity can be lower on coffee plants closer to *Inga* spp. as seen in our data as well (supplementary fig 1). Additionally, CBB infestation was lower in 2018 than 2019 (Supplementary fig 2) therefore it is possible that only on those plants with high ant activity, ants could locate and regulate the CBB in coffee fruits. In 2019, on the other hand, incidence of CBB in coffee fruits was significantly lower on plants with Azteca ants than on those with ant exclosures, irrespective of the shade tree treatment (Fig 6B). As stated earlier, CBB infestation was higher in 2019. When the density of CBB is high, the probability that ants will locate the berry borer is higher. Thus, even with lower ant activity in 2019 (ant activity was significantly lower only on plants paired with non-*Inga* spp.), ants were probably able to find and control the CBB more easily because they were present in higher densities.

We found no effects of proximity to *Inga* spp. on coffee floral traits and C:N or %N in coffee plants (supplementary figures 3 and 4). This is likely because all shade trees are pruned regularly and the leaf litter from both the N-fixing and non N-fixing shade trees gets mixed together on the ground. Additionally, coffee plants are also fertilized at the end of the flowering period using organic compost made on the farm. Thus, the mixing of the leaf litter of both N-fixing and non N-fixing shade trees along with fertilization using compost possibly masked the effects of enhanced N or other nutrients on coffee bushes paired with *Inga* spp. shade trees. While the direct effects of N-fixing shade trees were not apparent on plant traits or on resource availability, there was an indirect effect on ant activity which resulted in a difference in fruit weights and on the control of a major pest of *Coffea arabica*.

There was significant interannual variation in fruit quantity and the control of the coffee berry borer. Coffee plants had a significantly higher number of flowers in the second year, as well as higher number of visits by pollinators which probably resulted in a higher fruit set in the second year. Flowering in coffee plants is closely related to rainfall conditions preceded by a period of dry conditions (Peters and Carroll, 2012). Although we did not record abiotic conditions, it is possible that this change in flowering was probably due to interannual changes in rainfall in Chiapas, Mexico where the study took place. Finca Irlanda has also steadily increased the number of honeybee hives on the farm. Along with the simple increase in their abundance, mass flowering crops such as *Coffea arabica*, are more attractive to honeybees over other flowering resources (Bänsch et al., 2020) and together, this may have increased the number of visits in the second year compared to the first. The density of CBB increased in the second year while ant activity decreased, and these changes may be due to the natural oscillations that take place in predator-prey systems. Our study therefore emphasizes the need to study interactions among ES over a longer temporal period since interannual variation in the population sizes of organisms and interspecific interactions can alter the outcomes of ES interactions.

Our study highlights that ecosystem services are a result of a set of complex interactions and that management factors can have significant effects on both the provisioning of ecosystem services as well as on the interactions among them. It is therefore important to understand the conditions under which there might be synergies or trade-offs among ecosystem services if we want to manage agroecosystems both effectively and sustainably. Our study also highlights that shade trees are essential to the existence of *Azteca sericeasur*, and the service of pest control.

While the majority of the visits to coffee flowers were made by africanized honeybees, the most abundant bees in the region are all cavity nesting bees and shade trees are therefore essential for the service of pollination.

