## Supplementary Fig 1 for "Complex interspecific interactions influence the interactions between pest control and pollination in coffee agroecosystems"

Supplementary Information

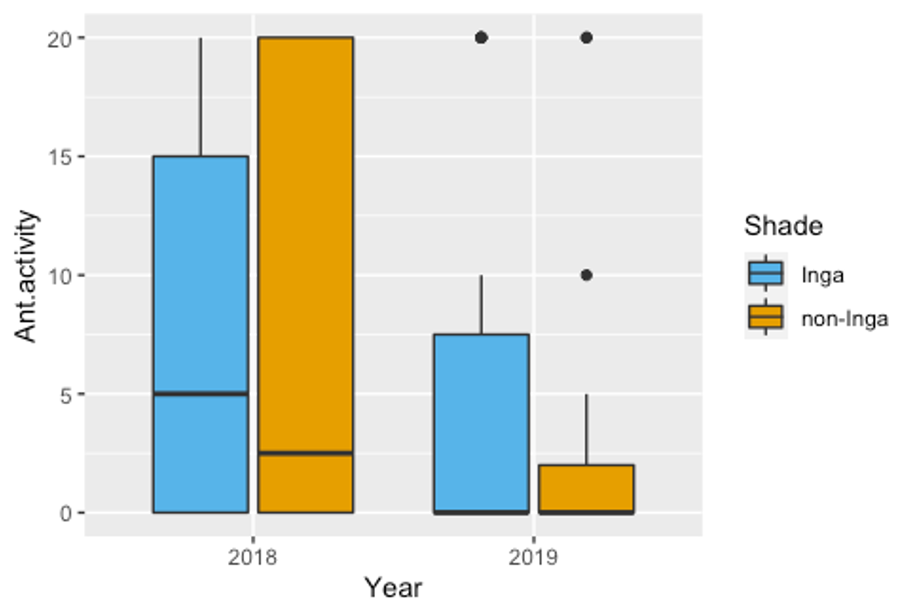

Supplementary Fig 1. Change in ant activity on shade trees from year 2018 to 2019.

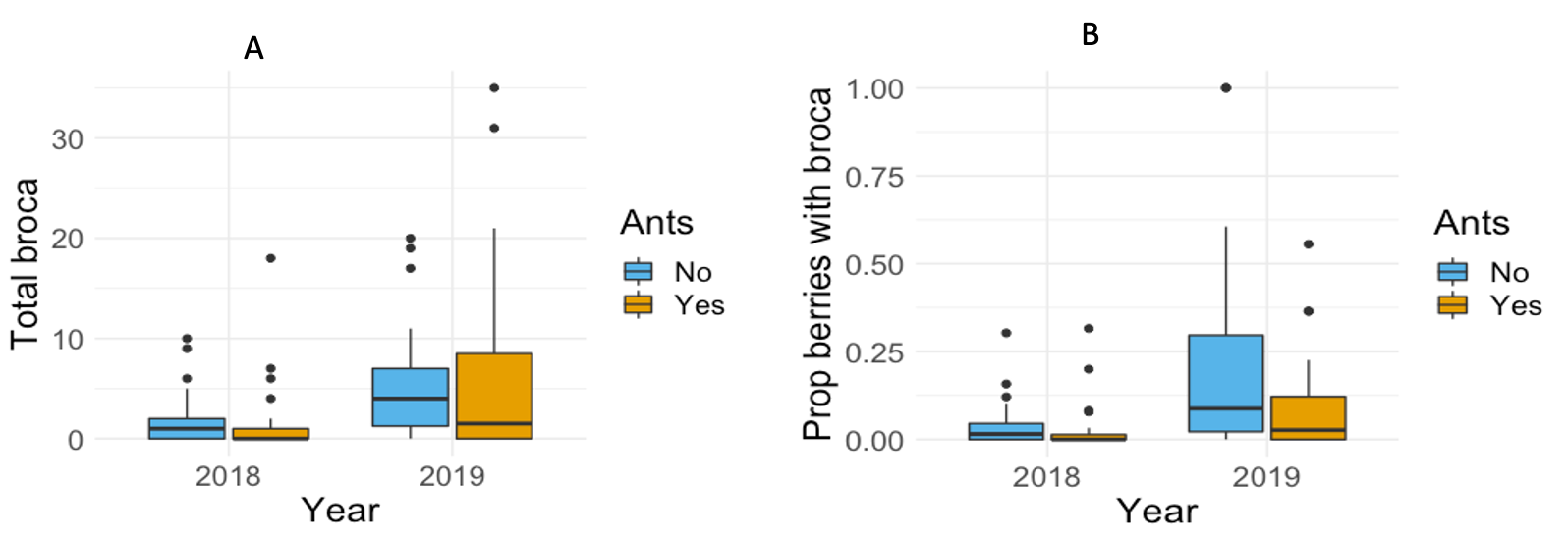

Supplementary figure 2 – A) Number of berry borers with and without ants in years 2018 and 2019. B) Proportion of berries with berry borer with and without ants in 2018 and 2019.

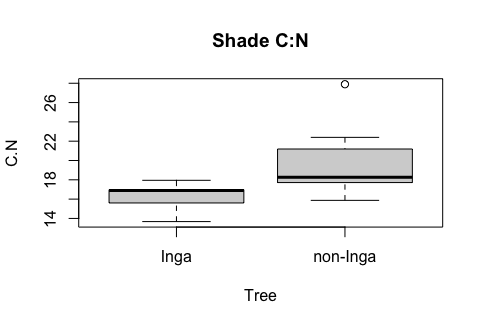

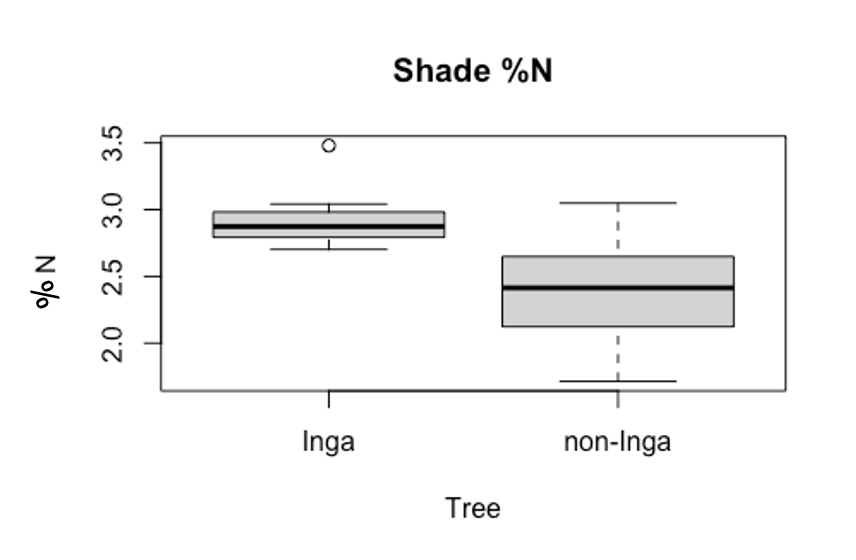

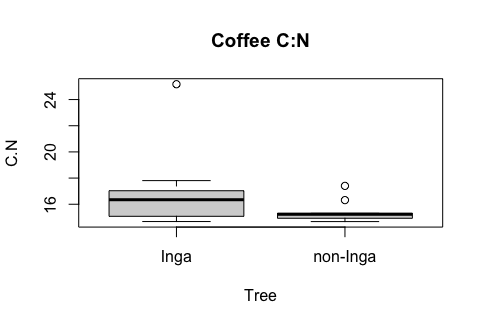

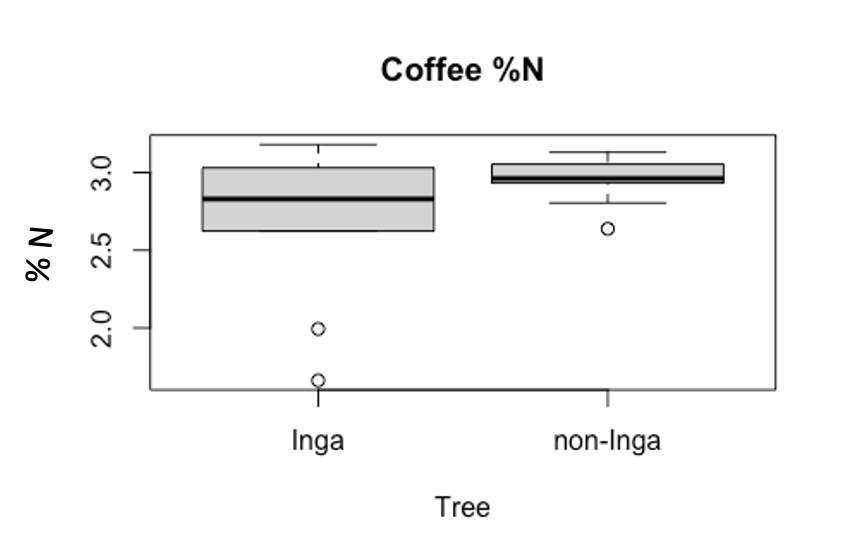

Supplementary Fig 3 Shade and coffee C:N ratios and %N

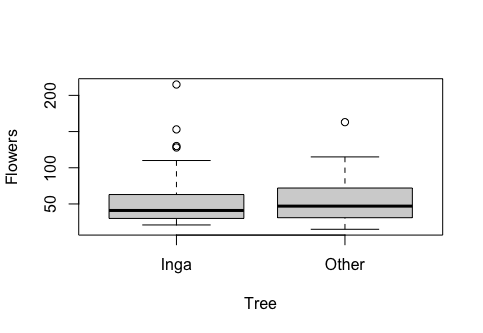

Supplementary Fig 4 Number of flowers on coffee bushes close to *Inga spp.* and non- *Inga spp.* shade trees.

Supplementary Table 1: Output of (generalized) mixed effects models with fruit weight as response variable and the number of beans and the presence/absence of CBB as explanatory variables. With each of bean, fruit weight increased by 0.57g and CBB presence decreased fruit weight by 0.11g.

|  | **WEIGHT** | | |
| --- | --- | --- | --- |
| *Predictors* | *Estimates* | *CI* | *p* |
| (Intercept) | 0.62 | 0.54 – 0.69 | **<0.001** |
| BEANS | 0.57 | 0.55 – 0.60 | **<0.001** |
| BROCA | -0.11 | -0.15 – -0.07 | **<0.001** |
| **Random Effects** | | | |
| σ^2^ | 0.13 | | |
| τ_00_ _Branch.id_ | 0.03 | | |
| τ_00_ _Tree.id_ | 0.03 | | |
| τ_00_ _Site_ | 0.00 | | |
| ICC | 0.34 | | |
| N _Site_ | 11 | | |
| N _Tree.id_ | 64 | | |
| N _Branch.id_ | 178 | | |
| Observations | 8083 | | |
| Marginal R^2^ / Conditional R^2^ | 0.173 / 0.453 | | |
